# Long-read sequencing reveals extensive *FMR1* somatic mosaicism in Fragile-X associated tremor/ataxia syndrome in human brain

**DOI:** 10.1101/2025.10.07.678589

**Authors:** Anna Dischler, Akshay Avvaru, Susana Lopez-Ignacio, Cristina Lau, Martin W. Breuss, Verónica Martínez Cerdeño, Harriet Dashnow, Caroline M. Dias

## Abstract

Fragile X–associated tremor/ataxia syndrome (FXTAS) is a progressive neurodegenerative disorder caused by a CGG repeat expansion in the 5’ untranslated region of the X-linked Fragile X messenger ribonucleoprotein 1 (*FMR1)*. Although the CGG repeat tract is known for instability that has been posited to contribute to clinical heterogeneity, the extent of somatic variation in human brain remains unclear, in part due to the technical limitations of sequencing long tandem repeats. Here, we quantified *FMR1* somatic variation in post-mortem brain tissue from individuals with FXTAS (n = 6) and Fragile X syndrome (FXS, n = 2) by applying amplification-free, targeted, long-read sequencing. This approach uncovered remarkable somatic mosaicism in repeat size and methylation in FXTAS, including somatic expansions and contractions which were not resolvable with traditional approaches. For example, in FXTAS, we identified somatic expansions to over 6000 base pairs in length as well as contractions to below the pathogenic range. We also identified unexpected patterns of methylation mosaicism on pre and full mutations. On the other hand, we replicated prior findings suggesting differential CGG expansion on the active X-chromosome in XX females. Finally, we examined the above cohort for expansions in 19 additional disease-associated repeat loci. Remarkably, we identified additional expansions in 5 out of 8 affected individuals, in *FXN* and *RFC1*. This work provides new insight into the extensive molecular variation underlying FXTAS in human brain and establishes a framework for studying repeat expansion disorders more broadly, highlighting the potential of long-read sequencing to advance our fundamental understanding of somatic mosaicism of these intractable regions of our genome.

## Introduction

Fragile X-Associated Tremor/Ataxia Syndrome (FXTAS) is an adult-onset neurodegenerative disorder that affects ∼10 – 60% of carriers of the X-linked *FMR1* premutation, a repetitive CGG sequence (∼55 – 200 repeats in length) [1-3]. The premutation is in the 5’ untranslated region of Fragile X Ribonucleoprotein 1 (*FMR1*) and is present at a population frequency of 1 in 400 males and 1 in 200 females, approximately [4-6]. Clinical features include ataxic gait, executive dysfunction, psychiatric symptoms, cognitive decline, neuropathy, with onset of neurological symptoms occurring approximately between ages 50 – 70 years [7-13].

Neuropathological signatures of this disorder indicate widespread brain dysfunction, including intranuclear inclusions in neurons and astrocytes, white matter degeneration, gliosis, and brain atrophy [14-16]. However, the severity and symptoms associated with FXTAS differ between individuals, and the source of this marked heterogeneity remains unknown.

An important feature of the premutation is that it is particularly unstable in both dividing and quiescent cells [17-20]. Multiple studies focusing on blood have demonstrated that length mosaicism, defined by varying lengths of the CGG repeat tract present within the same individual, and methylation mosaicism, when both methylated and unmethylated *FMR1* alleles are present, occurs frequently in individuals with the premutation and could drive clinical heterogeneity [21, 22]. The location of *FMR1*, the X-chromosome, could also theoretically influence instability [23, 24]. However, most studies are based on blood, and little is known about the extent of *FMR1* mosaicism across the brain.

Full mutations at *FMR1* exceeding 200 CGG repeats lead to Fragile X Syndrome (FXS), a neurodevelopmental disorder caused by *FMR1* hypermethylation and subsequent gene silencing, associated with intellectual disability, autism, and seizures [25-31]. Unlike FXS, premutation carriers display no hypermethylation, increased *FMR1* mRNA levels in blood, and variable levels of Fragile X Messenger Ribonucleoprotein (FMRP), the protein product of *FMR1* [32, 33]. The diverging clinical phenotypes implicate distinct molecular mechanisms in FXTAS pathophysiology, including gain-of-function effects, distinct from FMRP loss-of-function leading to synaptic dysfunction in FXS [34-37].

Although the repetitive nature of the CGG repeat has made it difficult for traditional tools to resolve the full extent of *FMR1* variability, recent developments in long-read technology have demonstrated increased sensitivity over repetitive genomic regions [38] and provide the additional benefit of simultaneous DNA methylation read-out. Targeted amplification-free approaches have also emerged for obtaining high coverage which is particularly useful for analyzing somatic mosaicism [38-42]. Here we applied long-read, high fidelity (HiFi) sequencing using a commercially available targeted amplification-free panel, PureTarget, to characterize *FMR1* length and methylation mosaicism (as well as at 19 additional disease-causing repeat loci across the genome) in human brain tissue from individuals with pre and full *FMR1* mutations.

## Results

### Determining FMR1 instability in a larger FXTAS cohort

Our initial FXTAS cohort consisted of 18 individuals (12 males, 6 females-Supplemental Table 1). We confirmed the genotype in all cases using triplet repeat primed PCR with capillary electrophoresis (TP-PCR/CE) on cerebellar hemisphere tissue (Figure 1a, Supplemental Figure 1a-r). Interestingly, full mutation expansions were detected in 22.2% of FXTAS cases (Supplemental Figure 1a, c, e, o). We then calculated the instability index (ii) of the *FMR1* premutation for all cases (Figure 1b). The ii (i.e. expansion index) is a metric that has been used to measure somatic length mosaicism of repeat expansion loci such as *FMR1, HTT* (huntingtin), and *FGF14* (fibroblast growth factor 14) using the electropherogram trace produced by TP-PCR/CE (Figure 1c) [23, 43-47]. We included the reference (modal) allele and all peaks to the right, that were a minimum of 20% relative fluorescence units (RFU) of the modal allele. Thus, an increased number of peaks above the threshold leads to a more positive index value. We excluded peaks to the left given these may include technical stutter peaks produced by short, incomplete PCR fragments [44, 47, 48]. Overall, we detected a broad range of ii values in both our male and female cohorts ranging from 0 – 7.7, allowing for distinction between cases with presumably low and high tendencies for somatic expansion/instability (Supplemental Figure 1s,t).

**Figure 1.**
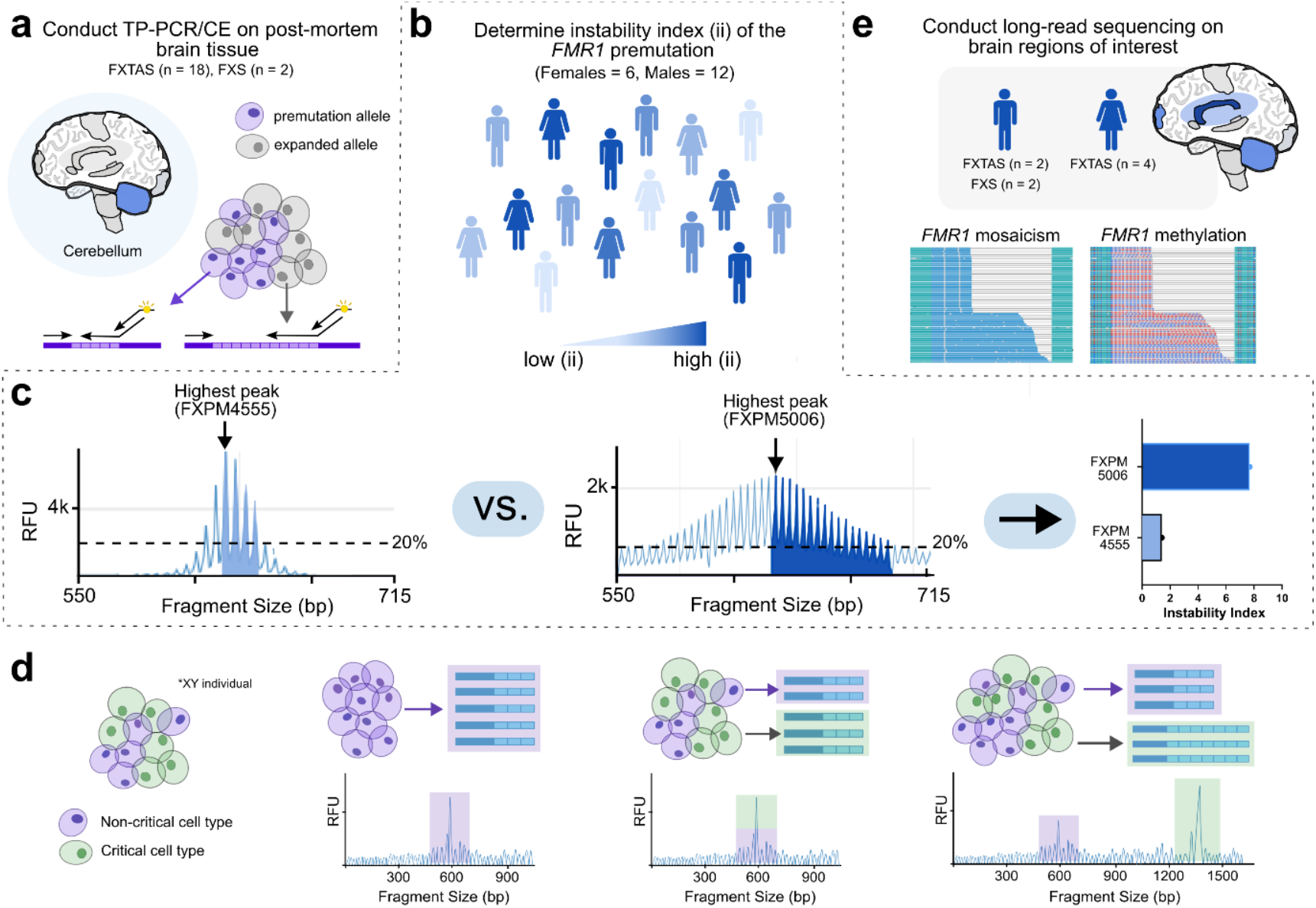
Characterization of *FMR1* somatic mosaicism in brain. a) TP-PCR/CE was conducted on cerebellar tissue from 18 individuals with FXTAS and two individuals with FXS to determine CGG repeat size. Cartoon demonstrates different length alleles captured from two theoretical cell populations. b) Instability index (ii) was calculated for 6 females and 12 males using electropherograms, with thresholding at 20% of the modal allele. c) Schematic demonstrates example of determining ii from samples FXPM 4555 and FXPM 5006. d) Schematic demonstrating how TP-PCR/CE would detect different theoretical examples of repeat expansion. Purple cells represent non-critical cell types, while green cells represent critical cell types. A single cell-type population with no mosaicism can be seen as a single peak on an electropherogram. A mixture of non-critical and critical cell-types may also have the same genotype (represented as a single peak). Over time, critical cell types may become burdened by extreme somatic expansion. e) Long-read sequencing was conducted on brain regions of interest from both males (n = 4) and females (n = 4) with either FXTAS (n = 6) or FXS (n = 2). HiFi sequencing and downstream analysis with TRGT detects mosaicism and methylation of *FMR1*, shown here in waterfall plots.

### Investigating FMR1 somatic length mosaicism with TP-PCR

Although instability values are helpful to determine case-to-case variability, it is unknown if patterns of *FMR1* instability throughout the brain are the same as the cerebellum alone. While the cerebellar hemisphere contains a dense layer of granule cells that predominate, other regions like the prefrontal cortex contain more heterogeneous cellular compositions. Given that it has been shown in other repeat expansion disorders like Huntington’s that certain critical cell types are disproportionately impacted by somatic expansions (Figure 1d) [49] it is possible there is significant regional variability in somatic mosaicism across the brain dependent on cell-type composition.

To screen for this, we conducted TP-PCR/CE on an additional 8 regions from case FXPM 5006 (ii = 7.7), the male FXTAS case with the highest ii (Supplemental Figure 1r). Each region exhibited a unique ii, with several regions like BA10 (ii = 10.6) and cerebellum (ii = 7.7) showing increased ii when compared to pons and BA4 (ii = 2.8, 6.8) (Supplemental Figure 2a,b). However, manual inspection of these traces suggested mosaicism across all regions (Supplemental Figure 2). We detected the presence of a full mutation in each region, although we could not accurately size the absolute length or cell fraction of the full mutation fragments with TP-PCR/CE given the known limitations of this approach (preferential amplification of smaller fragments). Interestingly, we found no significant correlation between ii values and either *FMR1* mRNA levels or FMRP expression within this case (Supplemental Figure 2c-f). This may relate to significant baseline differences in regional expression, as well as inability of the ii metric to accurately quantify variation seen in cases with significant mosaicism.

We also used TP-PCR to assess *FMR1* somatic mosaicism in a case with low a low ii, FXPM 4555 (ii = 1.4); however, we detected no regional variability across all five regions assessed (Supplemental Figure 3). Taken together, the above findings suggest that while *FMR1* length somatic mosaicism *is* observed across the brain, the extent of variability depends on the individual. Additionally, although TP-PCR may be sufficient for stable expansions, it does not fully capture somatic length mosaicism, including the absolute size or cell fraction of very long somatic expansions thus limiting its utility to quantify somatic mosaicism in cases with more marked variability.

### Long-read sequencing quality

We proceeded with long-read, amplification-free, targeted sequencing to better discern *FMR1* variability in 6 individuals with FXTAS and 2 with FXS with the PureTarget kit (Figure 1e). Coverage of *FMR1* was as high as 348x, and sequencing quality control analysis revealed a mean HiFi read length of ∼5 kilobases (Supplemental Data 1), consistent with the amplicon size of the targeted panel. Average Phred QV score of mapped reads was 44.6 (99.99684% accuracy), allowing for high confidence mapping of long repeat expansions (Supplemental Data 1). Although HiFi sequencing better covers through long repetitive regions, it is possible that the GC-rich premutation could still bias sequencing, as the coverage for *FMR1* was overall lower than other loci on the panel (Supplemental Data 1). To investigate this further we reviewed that coverage of premutation-length expansions in females with FXTAS vs their normal length alleles and found that coverage was comparable at ∼ 98x each on average (Supplemental Data 1), demonstrating little bias against repeats in the premutation range compared to normal length alleles. (Supplemental Data 1). Additionally, the female case and region with the highest *FMR1* coverage of all samples also had 8 reads in the full mutation range, including one read close to 3000 bp.Thus, even low frequency large expansions are captured with this approach.

### Long-read sequencing of FMR1 expansion in individuals with FXTAS and FXS

To observe patterns of somatic mosaicism of *FMR1* in our long-read sequencing, we proceeded with one female and one male FXTAS case, FXPM5006 and FXPM1008-20-RF, both of which demonstrated the highest ii out of their respective cohorts (Supplemental Figure 1s,t, Supplemental Table 2). We conducted bulk sequencing of cerebellum, periventricular white matter, and prefrontal cortex (BA10), as those regions are critical to FXTAS [50-60]. Post-mortem brain tissue from FXS, is less widely available. However, we were able to extend our analysis to two additional Fragile-X Syndrome cases, FXS 5319 and FXS 1031-09-LZ, to determine if full mutation *FMR1* repeat tracts >200 demonstrate length mosaicism patterns compared to the premutation alleles seen in individuals with FXTAS.

Sequencing of cerebellum, periventricular white matter, and BA10 of FXPM 5006 revealed *FMR1* premutation motif lengths ranging from 125 – 188, 108 – 197, and 57 – 198, respectively (Figure 2a-c). Interestingly, we detected a contracted, intermediate length, allele of 45 repeats in BA10 (Figure 2c) that had been undetected by TP-PCR. Reads supporting a full mutation expansion were identified in all three regions. In the cerebellum, there was only one read that supported a full mutation (2.2% total reads). However, expansions of over 1000 and 2000 repeats were detected in the periventricular white matter and BA10, respectively, with full mutation reads representing 14.6% and 33.3% of total reads for those regions. The largest range of repeat motifs was detected in BA10 (Fig. 2d).

Sequencing of the cerebellum, periventricular white matter, and BA10 of female case, FXPM 1008-20-RF, revealed premutation motif lengths ranging from 87 – 121, 86 – 125, and 71 – 195, respectively (Fig. 2e-g). We detected eight full mutation reads in the BA10 that ranged from 202 – 2832 motifs, equating to 4.57% of mutation-length reads (Fig. 2g). We did not detect full mutations in any other region. Again, the region with the largest range of *FMR1* mutation-length alleles was BA10 (Fig. 2h).

Sequencing of the cerebellum, periventricular white matter, and BA10 in FXS 5319, captured a range of full mutation variants with repeat sizes ranging from 300 – 387, 249 – 486, and 255 – 434, respectively (Fig. 2i-k) with the periventricular white matter containing the largest range of *FMR1* alleles (Fig. 2l). Additionally, only full mutation reads were observed in case FXS 1031-09-LZ (see *Methylation patterns* below). Interestingly, we did not detect any extremely long expansions or contractions like in the FXTAS cases in either FXS case. Overall, length distribution of *FMR1* varied from region to region, but higher magnitudes of somatic variation were observed in FXTAS cases rather than FXS.

**Figure 2.**
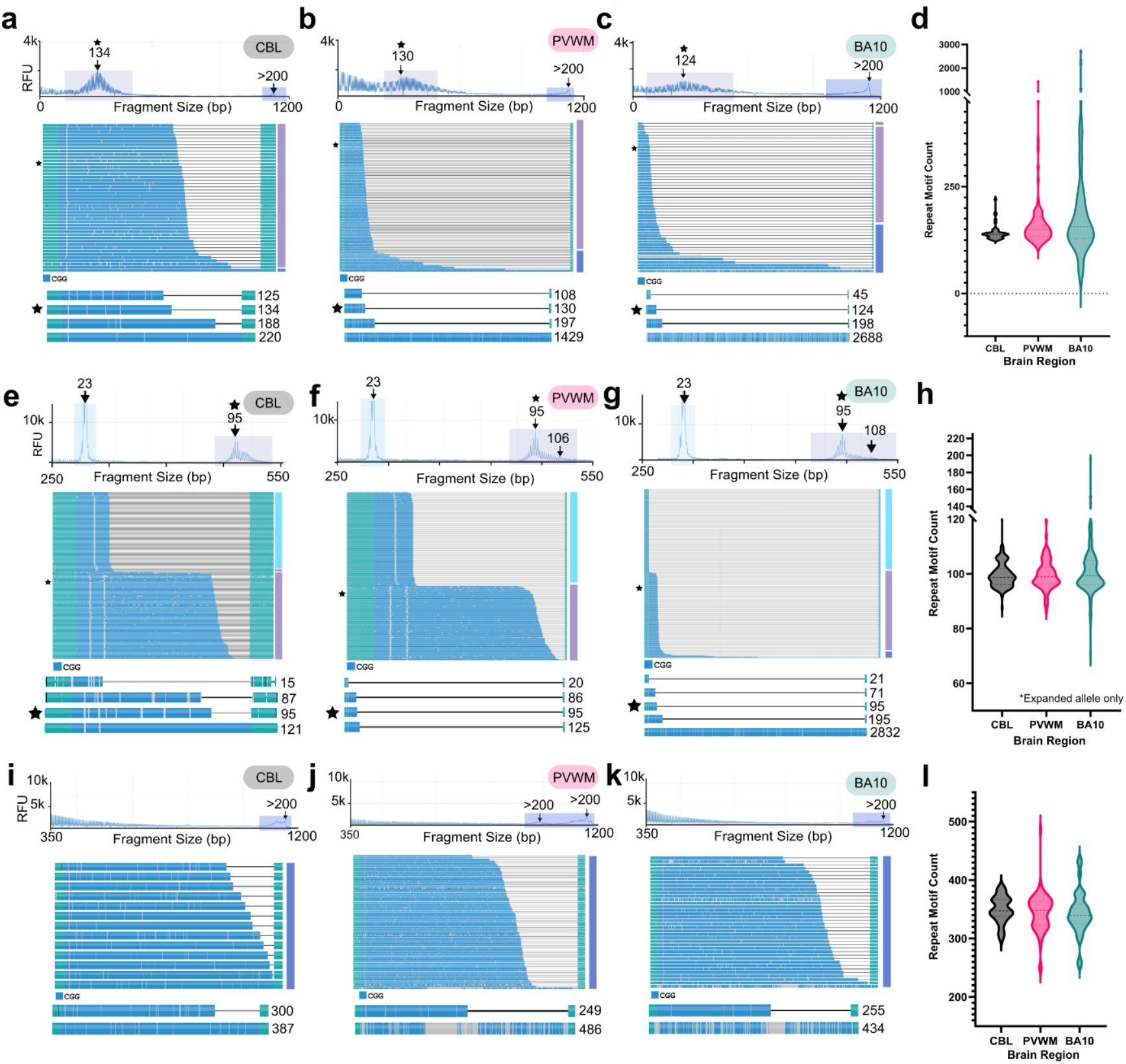
Targeted, long-read sequencing reveals variability in *FMR1* length mosaicism in human brain. a-c) TP-PCR/CE (top electropherograms) and long read sequencing (waterfall plots) from FXPM 5006 using cerebellum (a), periventricular white matter (b), or BA10 (c). Below each waterfall plot is shortest, modal (annotated with star), longest premutation, and longest full mutation number of repeats in descending order. d) Violin plot shows distribution of *FMR1* motif size by brain region in case FXPM 5006.e-g) TP-PCR/CE (top electropherogram) and long read sequencing (waterfall plots) from FXPM 1008-20-RF (female) using cerebellum (e), periventricular white matter (f), or BA10 (g). Below each waterfall plot is shortest normal length repeat, shortest premutation, modal premutation and longest full mutation (if applicable) number of repeats in descending order. h) Violin plots depict *FMR1* motif size by brain region for case FXPM 1008-20-RF. i-k) TP-PCR/CE (top electropherogram) and long read sequencing (waterfall plots) from FXS 5319 using cerebellum (i), periventricular white matter (j), or BA10 (k). Below each waterfall plot the shortest and longest motif in descending order. Only full mutation motifs over >200 were detected in all three regions for this case. l) Violin plots showing distribution of *FMR1* motif size by brain region for FXS 5319.

### Methylation patterns of FMR1

The amplification-free sequencing approach used here preserves methylation, allowing inference of 5-methylcytosine methylation data at CpG sites from enzyme kinetics used to generate the HiFi reads. We used this opportunity to analyze patterns of *FMR1* methylation in the FXTAS and FXS cases above on both a per cytosine and per read basis (See *Methods-Bioinformatics*).

While methylation analysis of male FXTAS case (FXPM 5006) revealed most premutation repeats were unmethylated as expected, we detected a small percentage of reads that we considered fully methylated, equating to 2.04% of reads in the cerebellum, 5.88% in the periventricular white matter, and 6.38% in BA10; this was not related to base call quality (Figure 3a-c, Supplemental Figure 4). We also noticed that several full mutation reads remained unmethylated, contrary to expectations. To corroborate these findings, we sequenced an additional male FXTAS case with known full mutation mosaicism as detected by TP-PCR, FXPM 5746 (Supplemental Figure 1e). We captured both the premutation and full mutation reads, but again, full mutation reads remained unmethylated (Supplemental Figure 5a-c). Thus, somatic length mosaicism to the full mutation range may be associated with unique methylation signatures in brain in FXTAS.

**Figure 3.**
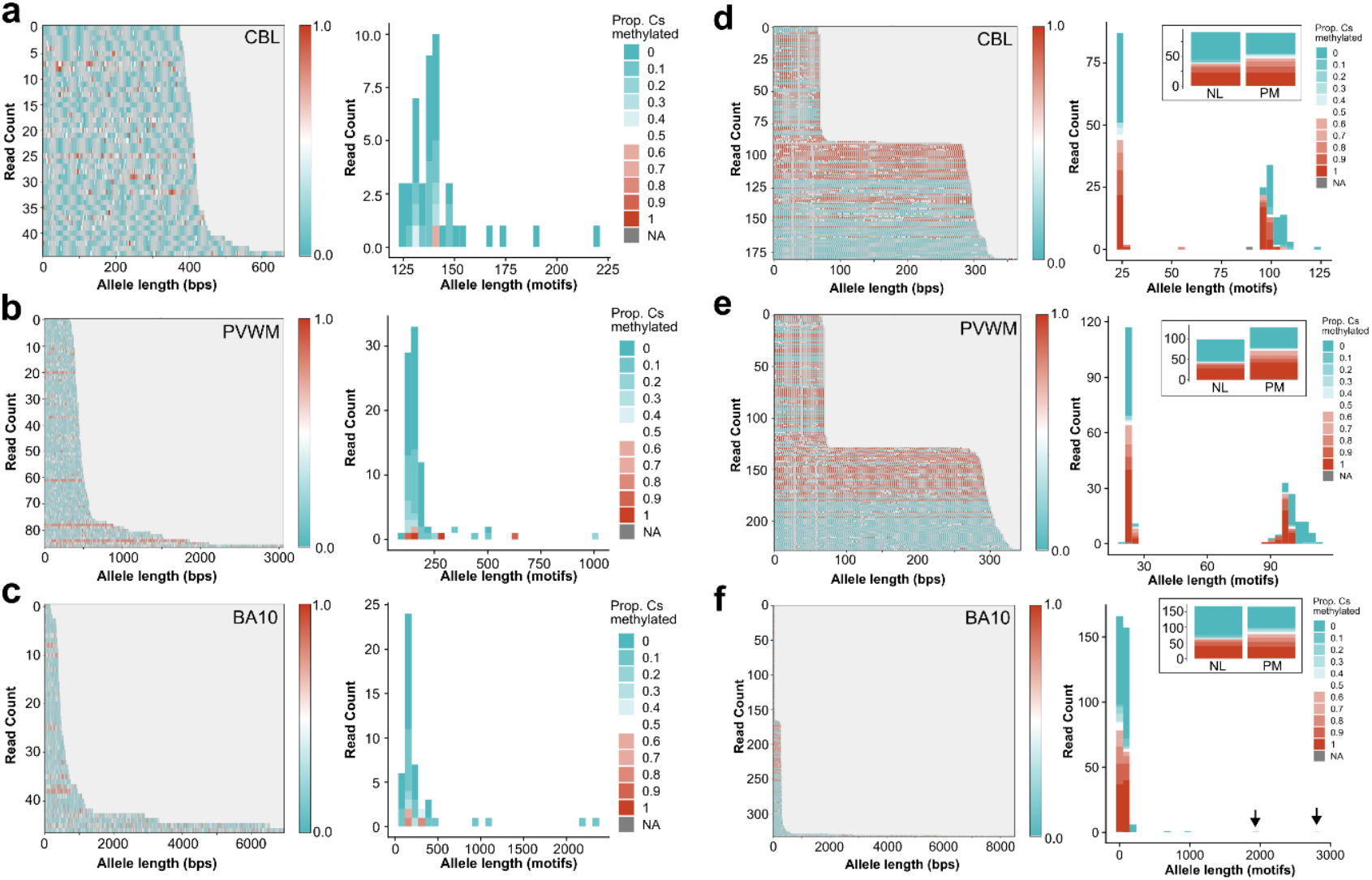
Variable methylation patterns of *FMR1*. a-c) Methylation analysis of FXTAS case FXPM 5006 depicting cerebellum (CBL) (a), periventricular white matter (PVWM) (b), and BA10 (c). Methylation waterfall plots demonstrate per-cytosine methylation probability (0 = unmethylated to 1 = methylated). Allele distribution plots demonstrate the distribution of reads by length, with color code indicating proportion of cytosine methylated along a read (0 = unmethylated, 1 = 100% cytosines >=0.66 (methylated), NA = no methylation information). a) Methylation was observed in only one premutation read. All full mutation reads remained unmethylated in CBL. b) Methylation waterfall plot reveals methylation in both premutation, and full mutation reads in PVWM. Allele distribution plot depicts both methylated and unmethylated pre and full mutations. c) Some premutation reads demonstrated a high proportion of methylated cytosines, while multiple full mutation reads demonstrated no methylation. Allele distribution plot depicts several unmethylated full mutations >1000 repeats in BA10. **d-f)** Intra-individual analysis of case FXPM 1008-20-RF in CBL (d), PVWM (e), and BA10 (f). Insets show methylation data of all reads supporting either the normal or premutation allele for that region. d) Methylation was observed on some normal and premutation reads. Allele distribution plot depicts peaks at ∼25 and ∼100 repeats. e) Methylation was observed on some normal and premutation reads. Allele distribution plot depicts peaks at ∼25 and ∼100 repeats. f) Methylation waterfall plot depicts methylation of normal and premutation alleles, but not the full mutation alleles.

In females, the correct dosage of X-linked genes is maintained through X chromosome inactivation (X-inactivation) [61-64]. Therefore, methylation along a read may also reflect patterns of X-driven, stochastic silencing caused by X-inactivation. In the previous female case (FXPM 1008-20-RF), we observed expected methylation on a portion of normal and premutation length reads (Fig. 3d-f), but also detected eight full-mutation expansions in BA10 that remained unmethylated and ranged from 202 – 2832 repeats (Fig. 3f). Interestingly, we noted that longer premutation alleles were frequently unmethylated (Fig. 3d-f, Supplemental Figure 4d-f) as has been previously reported. This finding held true when sequencing additional female cases (Supplemental Figure 5d-i).

Analysis of FXS 5319 demonstrated extensive methylation of the full mutation *FMR1*, with little variability; we did not detect major somatic mosaicism methylation differences in the regions assessed (Supplemental Figure 6a-i). Sequencing an additional FXS case, FXS 1031-09-LZ, also revealed expected hypermethylation of the locus with minimal variability (Supplemental Figure 7a-c). In summary, these results reveal unexpected somatic methylation patterns. Evidence of methylated premutation repeats, and full mutation repeats lacking methylation in both males and females with FXTAS counters dogma that expansion of the repeat to >200 repeats is necessary to trigger hypermethylation in the context of somatic mosaicism.

### Variation at repeat expansion disease loci outside of FMR1

Repeat expansion disorders, many of which disproportionately impact the nervous system, affect as many as 1 in 283 (0.35%) of people worldwide. For example, expansions in replication factor C subunit 1 (*RFC1*) and frataxin (*FXN*) are also associated with neurologic disorders [65-67]. *FMR1* expansions have previously been associated with disruptions in higher level chromatin organization that may contribute to genome-wide STR instability [68-71]. However, co-occurrence of expansions within an individual clinically may be underappreciated given historically, individual methods are needed to accurately detect each loci. The panel used here (PureTarget) detects 19 additional disease-associated repeat expansion loci (Supplemental Data), powering us to discover rare expansions at these additional loci in our cohort with a high confidence approach.

We sequenced a total of 14 regions across 8 affected individuals (FXTAS = 6, FXS = 2) and discovered additional germline expansions in 5 individuals (FXTAS = 3, FXS = 2).

Interestingly, 4 out of 5 expansions were in *RFC1*, with similar, non-pathogenic biallelic AAAAG_12-200_ expansions in two FXTAS cases (FXPM 5746 and FXPM 1030-09-MM). We noted that somatic expansion was present on both alleles; however, the degree of expansion differed by allele (Fig. 4a, b). Surprisingly, both FXS cases sequenced had uncommon, heterozygous expansions in *RFC1*. One monoallelic expansion consisted of ∼30 repeats of an AAAGGG hexanucleotide motif (Fig. 4c), while the other contained ∼60 repeats of an AAGAG pentanucleotide motif (Fig. 4d). Some minimal length mosaicism was observed on both.

One individual (FXPM 1008-20-RF) was a carrier for a pathogenic GAA_60+_ expansion in *FXN*. This locus is complex and may contain mononucleotide adenine repeat insertions within the GAA motif itself, as well as TTC repeat insertions. Due to this, size of the GAA repeat is difficult to discern. However, long-read sequencing captured a GAA expansion in cerebellum, periventricular white matter, and BA10, with the largest expansion in each region equating to 7189 bp, 5199 bp, and 3500 bp, respectively (Fig. 4e). The cerebellum demonstrated the largest range of expanded alleles, equating to a difference of 5652 bp between the shortest and longest expansions, denoting significant somatic mosaicism. This pattern was also distinct from the pattern seen with *FMR1* length mosaicism observed in this same case (for which BA10 demonstrated the largest allelic range).

**Figure 4.**
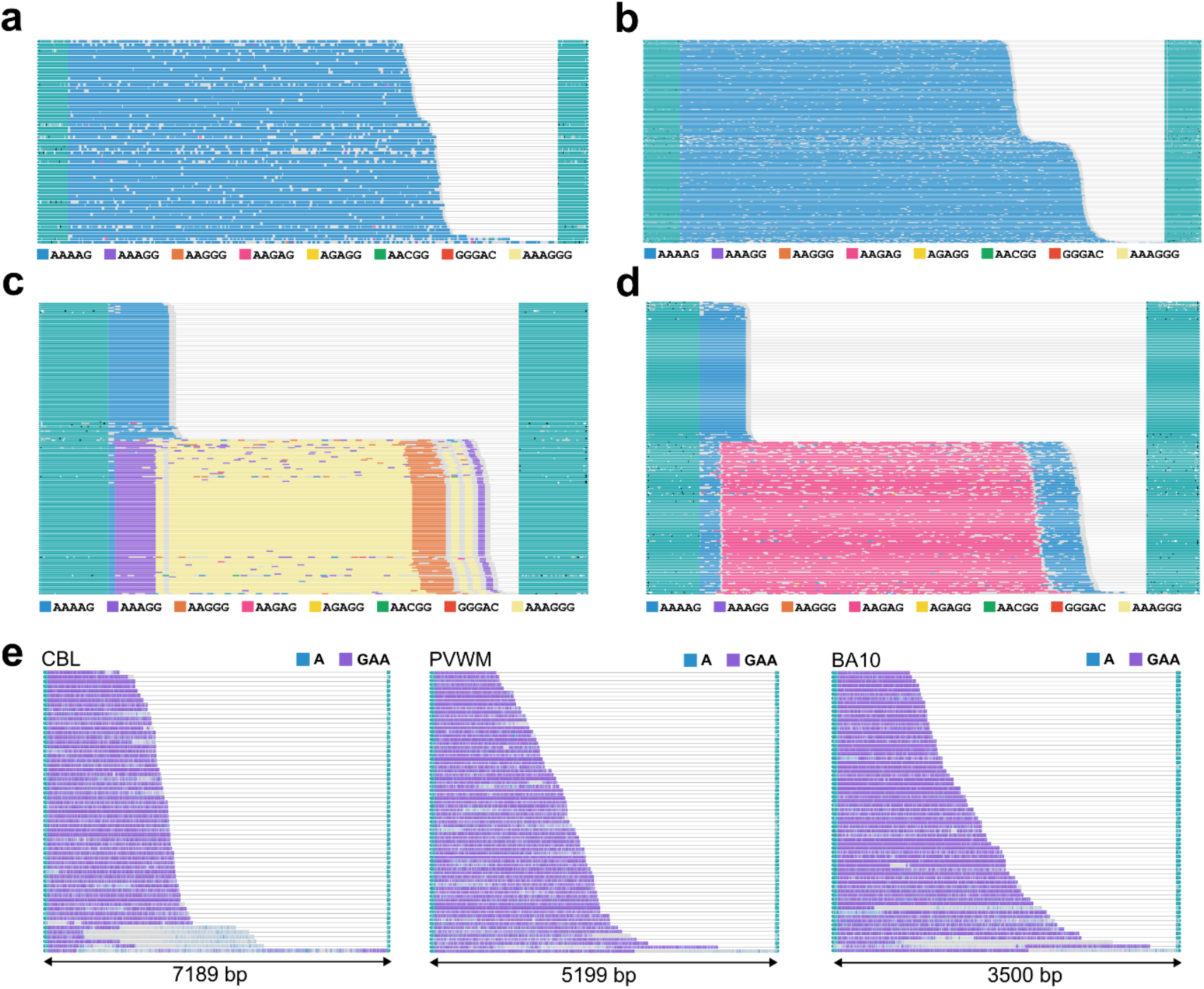
Somatic variation at repeat expansion disease loci outside of FMR1. a) Waterfall plots depict biallelic AAAAG expansion in *RFC1* in FXPM 5746. Modal motif size for each allele was 120 and 130 repeats. Somatic expansion of the motif was present on both alleles, expanding from the modal alleles by∼8 and ∼26 repeat motifs. b) Waterfall plots depict biallelic AAAAG expansion in *RFC1* in FXPM 1030-09-MM. Modal motif size for each allele was 95 and 115 repeats. Somatic expansion was present on both alleles, expanding from the modal alleles by ∼7 and ∼24 repeat motifs. c) Waterfall plots depict heterozygous AAAGGG expansion in *RFC1* in FXS 1031-09-LZ. d) Waterfall plots depict heterozygous AAGAG expansion in *RFC1* in FXS 5319. **e)** Waterfall plots depicting the expanded *FXN* allele in cerebellum (CBL), periventricular white matter (PVWM), and BA10 for FXPM 1008-20-RF. Length of smallest and largest inserts were 1537 bp and 7189 bp, 1142 bp and 5199 bp, and 1089 bp and 3500bp for CBL, PVWM, and BA10, respectively.

## Discussion

In this study we have demonstrated the advantage of using targeted amplification-free long-read sequencing to identify somatic variation of *FMR1* in brain from individuals with FXTAS and FXS, including expansions, contractions, and methylation mosaicism. Long-read sequencing provided accurate sizing of somatic length variation and subtle methylation mosaicism of *FMR1* beyond the detection capabilities of TP-PCR/CE.

Although TP-PCR/CE has become routine in both clinical and research settings [43, 44, 47, 48] our results suggest long-read sequencing is superior for interrogation of somatic mosaicism. Low frequency changes, including somatic expansions, contractions, and methylation alterations were frequently undetected with TP-PCR/CE that long-read sequencing easily distinguished. The ii values from TP-PCR/CE were important to determine case-to-case variation, as we successfully detected extreme expansions in cases with high ii values (Supplemental Figure 1, Fig. 2). However, regional differences within an individual could not be as robustly detected, which made characterization of somatic variation difficult to interpret via electropherogram (Supplemental Fig. 2a).

Our work does not investigate cell-type-specific mechanisms given limiting DNA input requirements for PureTarget. However, unlike the cerebellar hemisphere, which contains a high granule cell density, BA10 contains a more heterogeneous population including higher percentages of non-neuronal cell types. The cell-type-specific pathophysiology occurring in FXTAS [52] may influence premutation instability as has been observed in other repeat expansion disorders. However, future work will be required to answer this question more definitively, including whether critical cell types may drive length and/or methylation mosaicism. For example, on-going technical advances that reduce DNA library input requirements may allow for application of targeted long-read sequencing on fluorescent nuclear sorted brain cell populations in the near future.

Another critical aspect of interpreting molecular and clinical pathogenicity in Fragile X-related conditions involves knowledge of the methylation status of the expanded repeat region, particularly relevant to the phenomenon of X-chromosome inactivation in XX females. Sequencing of several female FXTAS cases revealed that longer premutation alleles within an individual are frequently unmethylated, corroborating prior work that shows *FMR1* premutation expansion predominantly occurs on the active X chromosome [23, 72, 73]. We should note that targeted HiFi reads also provided sufficient coverage to determine activation ratio, or the proportion of normal length allele on the active X chromosome. As we did not include any ‘gray zone’ carriers (∼41-54 repeats), we could not corroborate recent reports of hypermethylation in these alleles [74]. While we cannot exclude the possibility that hypermethylation represents an *FMR1* specific mechanism as opposed to reflecting X-chromosome inactivation, the lack of significant skew observed in our samples argues against this being the major driver in premutation methylation we observed in females. Given that activation ratio has been correlated with age of onset and FXTAS risk in women with the premutation [75-77], applying long-read sequencing in larger post-mortem brain cohorts in the future may illuminate fundamental drivers of cellular and clinical heterogeneity.

In Fragile X, traditionally, expansion of the CGGs to >200 repeats is associated with hypermethylation and silencing, presenting clinically as Fragile X-Syndrome. However, expansion alone is not sufficient to trigger hypermethylation of the repeat region [78-81], evidenced prominently here by the presence of unmethylated full mutations in multiple individuals with FXTAS. Unmethylated full mutations are not well described, with only a few case reports mentioning their detection [80, 82-85]. However, it is interesting that FXTAS somatic full mutations may not undergo silencing like FXS full mutations, suggesting unique regulatory mechanisms in the context of the premutation. For example, R-loops, secondary nucleic acid structures, are known to protect the promotor from methylation and impact transcription both in FXTAS and FXS; therefore, therapeutic approaches targeting R-loops have been undertaken [86-89]. Our work raises the question of how secondary structure formation may contribute to mosaicism in FXTAS [86]. Investigation using long-read technology is essential to better detect frequency of unmethylated full mutations in individuals with FXTAS and future studies are warranted to discern how these unexpected somatic patterns may contribute to FXTAS pathophysiology, heterogeneity, and ultimately therapeutic strategies, for example, to artificially contract expansions.

We identified additional germline expansions at loci outside of *FMR1*. The *FXN* expansion we detected was rare (population frequency of only 0.1 – 1.6% [90]) and demonstrated significant length mosaicism across brain regions; biallelic *FXN* expansions are causative of Friedrich’s ataxia [91]. The other expansions were detected in *RFC1*. Typically, biallelic AAGGG_250-2000_ expansions in intron 2 of *RFC1* cause Cerebellar Ataxia, Neuropathy and Vestibular Areflexia Syndrome (CANVAS), a disorder that affects sensory neurons, the vestibular apparatus, and the cerebellum [92, 93]. The expansions we detected are thus unlikely pathogenic; however, the AAAGGG hexanucleotide and AAGAG pentanucleotide motifs are extremely rare and detected at very low frequencies in gnomAD [94]. Given only limited flanking sequence is obtained, we are unable to make conclusions about expansion haplotypes across loci, however, it is interesting that1) we were able to detect additional rare expansions in 5 out of 8 individuals sequenced and 2) patterns of somatic length variation at *FMR1* and *FXN* differed within an individual. The latter suggests the possibility of distinct cellular mechanisms regulating this somatic variation for these two loci. The ability to resolve large, complex repeat expansions across multiple disease-associated loci beyond *FMR1* provides a unique opportunity to investigate co-occurrence of germline expansions as well as the shared and unique patterns of somatic variation of repeat expansion disorders within an individual in future work [95]. For example, do certain individuals demonstrate repeat hypermutability?

In summary, our work reveals extensive *FMR1* somatic mosaicism in Fragile-X associated tremor/ataxia syndrome in human brain. Overall, long-read sequencing provided a comprehensive view into patterns of this mosaicism suggesting the presence of extensive length and methylation mosaicism in some cases of FXTAS. While this cohort is too small to make any conclusions about the clinical significance of our findings, our work is consistent with prior reports of blood, demonstrating mosaicism is not restricted to hematologic lineages. Our findings suggest past reports dependent on TP-PCR approaches may have underestimated the true extent of somatic variation. For example, a single individual with FXTAS had full mutation reads ranging from 2-34%, depending on the brain region; these differences were not detectable via TP/PCR. Given mosaicism can be used as a tool to determine critical cell types relevant to pathophysiology, this variability has implications for understanding both basic science mechanisms, as well as clinical heterogeneity. The frequently observed full mutations that lack methylation provoke questions about differences in the endogenous cellular machinery that regulates repeat stability and methylation across development as compared to the germline. This is particularly critical to understand, given challenges in artificially contracting either full or premutation sized repeats to the “normal” range, an area of intense therapeutic interest. Additionally, ability to obtain high coverage data across repeat regions additional to *FMR1* may illuminate uncharacterized patterns of repeat instability across the genome in repeat expansion disorders.

## Methods

We used patient tissue from either the NIH NeuroBioBank or the UC Davis CENE tissue bank with approval from their institutional review boards after written informed consent was obtained. Initial dissection of frozen postmortem tissue for brain-bank specimens was done with sectioning under standardized procedures. Frozen tissue was stored at −80°C and kept frozen until processing.

Tissue (10 mg – 75 mg) was dissected from the larger tissue block, using pre-chilled sterile forceps and scalpels in a Leica cryostat CM1520 maintained at −20°C. All research on these deidentified specimens was performed at the University of Colorado Anschutz.

### DNA isolation

DNA isolation for TP-PCR/CE was completed using the DNeasy Blood and Tissue Kit (Qiagen, 69504) with ∼25 mg input tissue and with modifications to the manufacturer’s protocol, for postmortem brain tissue. DNA purity was measured using the NanoDrop 2000 (ThermoFisher) and DNA concentrations were obtained via the Qubit 2.0 Fluorometer and the dsDNA HS Assay kit (Invitrogen, Q32851).

### Triplet-Primed PCR and Capillary Electrophoresis

The *FMR1* PCR/CE Kit (Asuragen, 49402) was used throughout. DNA input ranged from 10-60 ng. After thermal cycling, samples were prepared with ROX1000 allelic ladder (Asuragen) and denatured at 95°C for two minutes with HiDi Formamide (Applied Biosystems, 4311320). Samples were analyzed on a SeqStudio Genetic Analyzer (Applied Biosystems, A35644). Electropherograms were analyzed using both AmplideX Reporter software (Asuragen) and GeneMarker (SoftGenetics).

### Quantitative PCR (qPCR)

Total RNA was extracted from 10 mg brain tissue using the RNeasy Micro kit (Qiagen, 74104) and reverse-transcribed with qScript Ultra SuperMix (Quantabio, 95217-100) according to the manufacturer’s instructions. Primer pairs for *FMR1, ACTB*, and *GAPDH* were designed to span exon–exon junctions and validated by standard curve analysis (5-point, 10-fold serial dilutions of pooled cDNA) (Supplemental Table 3). All assays demonstrated single-peak melt curves and no amplification in no-template controls. Relative expression values were calculated using efficiency-corrected ΔCt normalization [96], with normalization to the geometric mean of two reference genes (*ACTB* and *GAPDH*). Values are presented as normalized relative expression units. Each biological sample was run in triplicate technical reactions, which were averaged prior to analysis.

### Western Blotting

Protein lysates were prepared from postmortem brain tissue by homogenization by pestle using RIPA buffer (Thermo Fisher Scientific, 89901) supplemented with protease inhibitors (Roche, 11836170001). Lysates were cleared by centrifugation at 12,000 × g for 20 min at 4 °C, and protein concentrations were determined by BCA assay (Thermo Fisher Scientific, A55861). Protein was denatured in Laemmli buffer (Bio-Rad, 1610737), separated on 4–20% SDS-polyacrylamide gel (Bio-Rad, 4561093). Gel was transferred to nitrocellulose membranes (Bio-Rad, 1704158) and blocked for 1 h at room temperature in Intercept® (TBS) Blocking Buffer (LICORbio, 927-60001), then incubated overnight at 4 °C with primary antibodies against FMRP (Rabbit anti-FMRP, Cell Signaling Technologies, 4317S) at a 1:750 dilution and β-actin (Mouse anti-β-actin, Santa Cruz Biotechnology, sc-47778) at a 1:1,000 dilution. Membranes were washed with TBS-T (1x Tris Buffered Saline with 1% Tween 20) (Millipore Sigma T5912 and P9416) then incubated with a 1:10,000 dilution of conjugated secondary antibodies: IRDye® 800CW Donkey anti-Rabbit IgG and IRDye® 680RD Goat anti-Mouse IgG for 1 h at room temperature in the dark (LICORbio, 926-32213 and 926-68070). Blots were imaged on a LICOR Odyssey DLx imaging system and band intensities were quantified using Empiria Studio Software (LI-COR). Target protein signals were normalized to β-actin.

### Extracting high quality high molecular weight genomic DNA for PureTarget

High molecular weight DNA was extracted using the NanoBind PanDNA kit (PacBio, 103-260-000). Approximately 50 mg of tissue was finely minced with a scalpel and subjected to glass Dounce homogenization in 750 uL cold Buffer CT before proceeding to the extraction process. After, DNA was incubated at room temperature overnight. DNA purity and concentration was evaluated in triplicate from the upper, middle, and lower portion of the tube using a NanoDrop 2000 (ThermoFisher) and Qubit BR dsDNA assay (Invitrogen, Q32850) with the Qubit 2.0 Fluorometer. Each sample was tested for RNA contamination with the Qubit HS RNA assay (Invitrogen, Q32852). Size of DNA was measured on a 4200 TapeStation using gDNA tape to ensure 50% DNA > 30 kb.

### Generating PureTarget Repeat Expansion Libraries

DNA input for library preparation ranged from 0.5 ug – 4 ug. Library preparation was done using the PureTarget Repeat Expansion Panel kit as per kit instructions. Samples were pooled in batches of 8 or 16 and concentrated prior to HiFi sequencing on the Revio without SPRQ chemistry (PacBio).

### Statistics and Graphing

All statistical analyses and data visualization were performed in GraphPad Prism 10.3.0. Data are presented as mean ± standard error of the mean (SEM) unless otherwise noted.

### Bioinformatics

Raw data was imported from the Revio to SMRTLink v13.1 to generate HiFi reads using the PureTarget repeat expansion analysis workflow. This includes demultiplexing with lima and determining 5mC methylation probabilities for CpG sites. Reads are then mapped to the hg38 reference genome with pbmm2 1.14.0. Tandem repeat calling was conducted via TRGT 1.5.1 [97] with a min read quality of −1.0. TRGT input consisted of the hg38 reference genome, repeat expansion panel BED file, and pbmm2 mapped BAM file. Output consisted of a VCF file and spanning BAM file, both of which were sorted and indexed prior to visualization. Allele and waterfall plots, in addition to methylation plots, were created with the TRGT --plot function using either TRGT 1.5.1 or TRGT 3.0.0. Graphpad Prism 10 was used for statistics and plotting.

We developed an optimized method to extract per-base sequence and methylation metrics from each read, tailored for targeted long-read sequencing of tandem repeat loci. The script retrieves reads that map to the target repeat locus. For each read, it identifies the upstream and downstream flanking sequences of the repeat locus, then extracts the bases located between these flanks, corresponding to the repeat region. The allele length of the repeat is defined as the length of the sequence between the flanks. The average base-call quality across the repeat region is calculated for each read. The code implementation of the method is available as a GitHub repository: https://github.com/dashnowlab/FMR1-CGG-instability.

Per-base methylation probability scores are reported on a scale of 0 to 1 in the alignment file, where 0 = unmethylated and 1 = highest probability methylated. To simplify interpretation, residues with a methylation score >= 0.66 were classified as methylated, while those with score of<=0.33 were classified as unmethylated. Then, proportion of methylated cytosines is calculated and summed up on a per-read basis. For example, a read containing 10 CpGs, with 8 CpGs scoring >= 0.66 signifies the overall methylation proportion score for that read is 0.8. These data are then used to investigate the relationship between allele length and methylation patterns across individual reads.

## Acknowledgements

We thank the individual donors and their families, as well as the NIH NeuroBioBank (including The University of Maryland Brain and Tissue Bank, and The Human Brain and Spinal Fluid Resource Center -Sepulveda) as well as the UC Davis CENE brain bank for providing tissue for these studies. High molecular weight DNA sizing and QC was conducted by the Genomics Shared Resource at University of Colorado - Anschutz Medical Campus (Research Resource Identifier RRID: SCR_021984). Capillary electrophoresis was conducted by the Organoid and Tissue Modeling Shared Resource at University of Colorado - Anschutz Medical Campus (RRID: SCR_021993). We thank the Arizona Genomics Institute at the University of Arizona, Arizona for assistance with PacBio sequencing. We thank Egor Dolzhenko for helpful discussions around bioinformatic analyses and Tamim Shaikh for thoughtful comments on the manuscript.

## Funding

CD is supported by the Boettcher Foundation (Webb-Waring Biomedical Research Award). SL was supported in part by the Randi J. Hagerman Summer Scholar Research Award from the National Fragile X Foundation. AA and HD are supported by NHGRI grant 4R00HG012796-03 and NHMRC Investigator grant GNT2026126. VMC is supported by NS107131. RRID: SCR_021984 is supported by the Cancer Center Support Grant P30CA046934. RRID: SCR_021993 is supported by NIH grants P30CA046934, P30DK116073, and the CU Anschutz Gates Institute and Section of Developmental Biology.

## Conflict of Interests

The authors have no conflicts of interest to disclose.

## Abbreviations

AGG: Adenine-guanine-guanine
BA10: Brodmann Area 10
CANVAS: Cerebellar Ataxia, Neuropathy, and Parkinson’s Disease
CBL: Cerebellum
CC: corpus callosum
CGG: Cytosine-guanine-guanine
FGF14: Fibroblast growth factor 14
FMR1: Fragile X messenger ribonucleoprotein 1
FMRP: Fragile X Messenger Ribonucleoprotein
FXN: Frataxin
FXPM: Fragile-X permutation
FXS: Fragile-X Syndrome
FXTAS: Fragile-X Tremor/Ataxia Syndrome
HTT: Huntingtin
ii: Instability index
PVWM: Periventricular white matter
PCR: Polymerase chain reaction
RFC1: Replication Factor C Subunit 1
RFU: Relative fluorescence units
TP-PCR/CE: Triplet primed polymerase chain reaction with capillary electrophoresis
UFM: Unmethylated full mutation

## Author Contributions

**Conceptualization:** Anna Dischler, Caroline Dias **Methodology** Anna Dischler, Caroline Dias, Akshay Avvaru, Harriet Dashnow **Software/Programming** Akshay Avvaru, Anna Dischler, Harriet Dashnow **Validation** Anna Dischler, Caroline Dias **Formal/Bioinformatic Analysis** Anna Dischler, Akshay Avvaru, Harriet Dashnow **Investigation** Anna Dischler, Susana Lopez-Ignacio, Cristina Lau **Writing - original draft preparation:** Anna Dischler, Caroline Dias **Visualization:** Anna Dischler, Akshay Avvaru **Supervision:** Caroline Dias, Martin Breuss, Harriet Dashnow **Resources:** Verónica Martínez Cerdeño, Harriet Dashnow All authors contributed to the review, editing, and final approval of the manuscript.

## Data Availability

All sequencing data is available through the controlled access section of dbGaP, accession # pending. Code implementation of methods used are available on Github: https://github.com/dashnowlab/FMR1-CGG-instability.

## Notes

### Competing Interest Statement

The authors have declared no competing interest.

