## Supplement for "Long-read sequencing reveals extensive *FMR1* somatic mosaicism in Fragile-X associated tremor/ataxia syndrome in human brain"

### Supplemental Table 1: Donor demographics and characterization

| Identifier | Diagnosis | Age | Sex | Race | PMI | RIN | TP-PCR |
|---|---|---|---|---|---|---|---|
| FXPM5006 | FXTAS | 85 | M | White | 5 | 7.1 | CBL, CC, BA10, BA17, BA4, BA42, PVWM, NA, Pons |
| FXPM5746 | FXTAS | 80 | M | White | 22 | Unknown | CBL, BA22 |
| FXPM4555 | FXTAS | 80 | M | White | 12 | Unknown | CBL, BA10, NA, CC, PVWM |
| FXPM1011-05-RD | FXTAS | 82 | M | Unknown | Unknown | Unknown | CBL |
| FXPM1006-14-JG | FXTAS | 81 | M | Unknown | 11 | Unknown | CBL |
| FXPM1002-04-ER | FXTAS | 87 | M | Unknown | Unknown | Unknown | CBL |
| FXPM334-03-KC | FXTAS | 81 | M | Unknown | Unknown | Unknown | CBL |
| FXPM1007-04-LL | FXTAS | 82 | M | Unknown | Unknown | Unknown | CBL |
| FXPM1018-08-EB | FXTAS | 79 | M | Unknown | 7.25 | Unknown | CBL |
| FXPM1061-08-AM | FXTAS | 82 | M | Unknown | 29 | Unknown | CBL |
| FXPM1004-10-MG | FXTAS | 77 | M | Unknown | 5.25 | Unknown | CBL |
| FXPM1009-18-JF | FXTAS | 72 | M | Unknown | 18 | Unknown | CBL |
| FXPM1006-06-CM | FXTAS | 80 | F | Unknown | 5 | Unknown | CBL |
| FXPM1004-019-EA | FXTAS | 89+ | F | Unknown | 4 | Unknown | CBL |
| FXPM1041-08-IM | FXTAS | 79 | F | Unknown | Unknown | Unknown | CBL, CC |
| FXPM1015-09-DK | FXTAS | 89+ | F | Unknown | Unknown | Unknown | CBL |
| FXPM1008-20-RF | FXTAS | 80 | F | Unknown | 21 | Unknown | CBL, PVWM, BA10 |
| FXS5319 | FXS | 71 | M | White | 17 | 6.6 | CBL, BA10, PVWM |
| FXS1031-09-LZ | FXS | 64 | M | Unknown | 11.5 | Unknown | CBL |

### Supplemental Table 2: Long-read sequencing of affected individuals

| | Identifier | Diagnosis | PureTarget |
|---|---|---|---|
| Intra-individual sequencing | FXPM5006 | FXTAS | CBL, PVWM, BA10 |
| | FXPM1008-20-RF | FXTAS | CBL, PVWM, BA10 |
| | FXS5319 | FXS | CBL, BA10, PVWM |
| | **Identifier** | **Diagnosis** | **PureTarget** |
| Inter-individual sequencing | FXPM5746 | FXTAS | BA22 |
| | FXPM1041-08-IM | FXTAS | CC |
| | FXPM1015-09-DK | FXTAS | CBL |
| | FXPM1033-09-MM | FXTAS | CBL |
| | FXS1031-09-LZ | FXS | CBL |

### Supplemental Table 3: qPCR primers

| Primer # | Primer Name | Sequence |
|---|---|---|
| 1 | hACTBexon4-5 F | GGACTTCGAGCAAGAGATGG |
| 2 | hACTBexon4-5 R | AGCACTGTGTTGGCGTACAG |
| 3 | hGAPDHexon3-4 F | TCACCAGGGCTGCTTTTAAC |
| 4 | hGAPDHexon3-4 R | TGACGGTGCCATGGAATTTG |
| 5 | hFMR1exon8-9 F | CCTGGGGTCACTGCTATTGA |
| 6 | hFMR1exon8-9 R | TCTTCAGCAAATTCGAGAAAGCT |

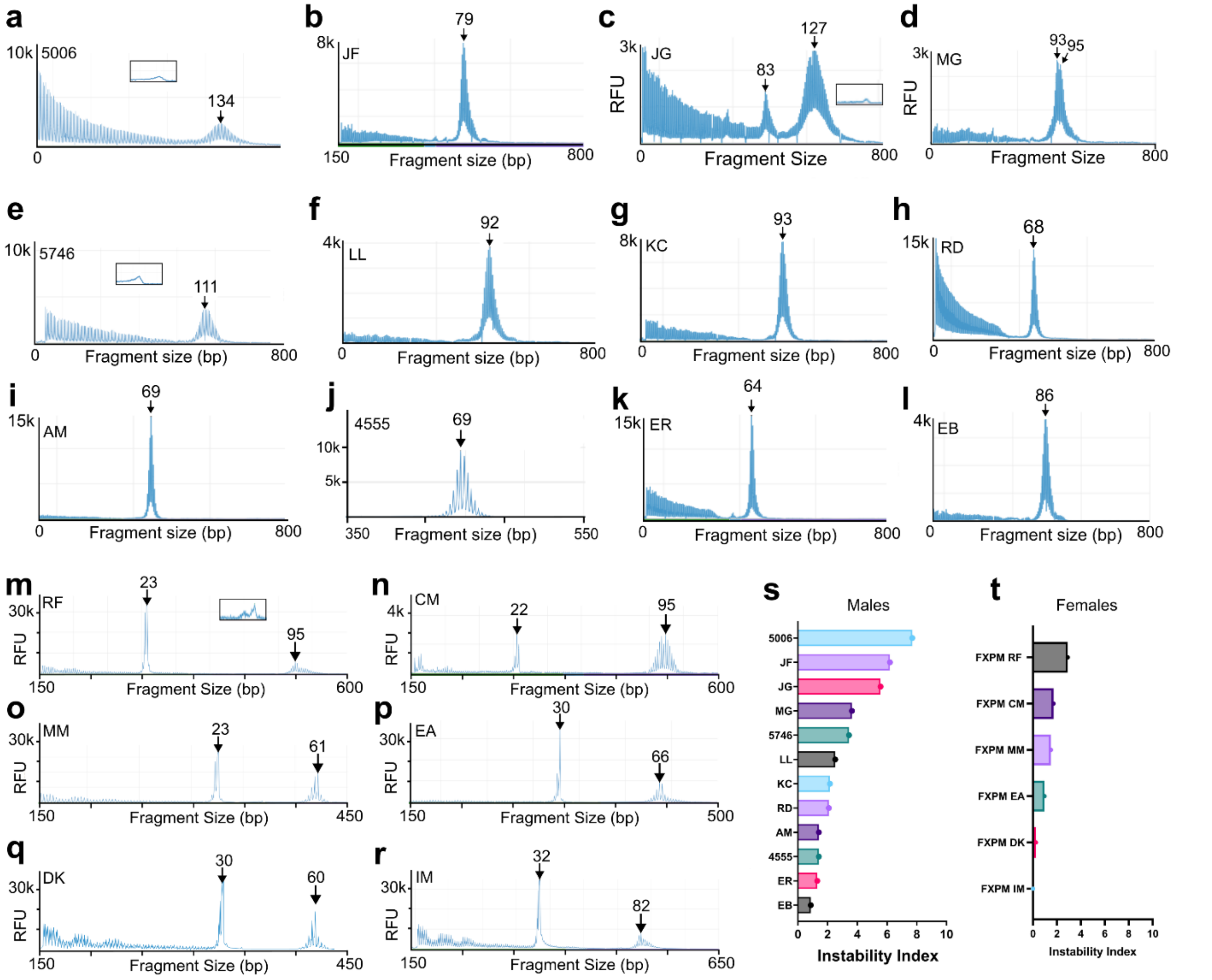


**Supplemental Figure 1. *FMR1* patterns of instability in larger a FXTAS cohort. a-l)** Cerebellar TP-PCR/CE of male FXTAS cohort in order of instability index value: *5006* (a), 1009-18-*JF* (b), 1006-14-*JG* (c), 1004-10-*MG* (d), *5746* (e), 1007-04-*LL* (f), 334-03-*KC* (g), 1011-15-*RD* (h), 1081-08-*AM* (i), *4555* (j), 1002-04-*ER* (k), 1018-08-*EB* (l). **m-r)** Cerebellar TP-PCR/CE of female FXTAS cohort in order of instability index value: 1008-20-*RF* (m), 1006-06-*CM* (n), 1030-09-*MM* (o), 1004-019-*EA* (p), 1015-09-*DK* (q), 1041-08-*IM* (r). s) Instability indices of male FXTAS cohort. t) Instability indices of female FXTAS cohort. Insets show zoomed in full mutation expansions > 200 bp.

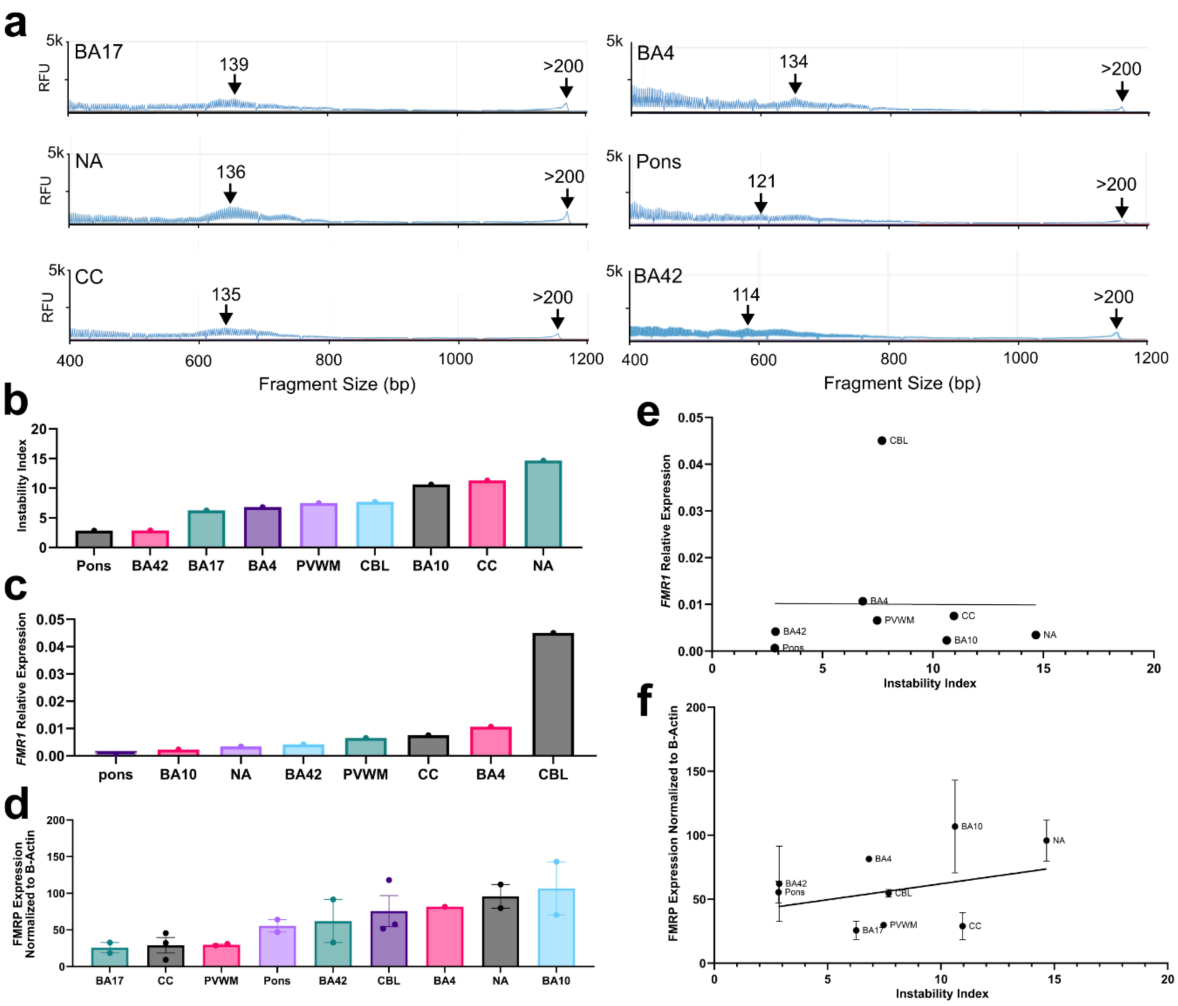


**Supplemental Figure 2. *FMR1* TP-PCR in high (ii) individual.** Molecular characterization from case FXPM 5006. Regions assessed: pons, BA42, BA17, BA4, periventricular white matter, cerebellum, BA10, corpus callosum, and nucleus accumbens. a) Electropherograms from brain region specific TP-PCR/CE depicting premutation allele at 139 repeats in BA17, 136 in nucleus accumbens, 135 in corpus callosum, 134 in BA4, 121 in pons, and 114 in BA42. Electropherograms for cerebellum, BA10, and periventricular white matter excluded. b) Instability index calculations. c) Bar plot depicting relative expression of *FMR1* determined via qPCR (relative expression = *ACTB* and *GAPDH*). Excludes BA17 due to poor RNA quality. d) Bar plot depicting FMRP expression normalized to Beta Actin determined by Western blot. e) Linear regression of *FMR1* relative expression against instability index across all brain regions ($\beta = -2.6x10^{-5}$ , $R^2 = 5.1x10^{-5}$, $p = 0.99$). f) Linear regression of FMRP expression against instability index across all brain regions ($\beta = 2.5$, $R^2 = 0.07$, $p = 0.27$).

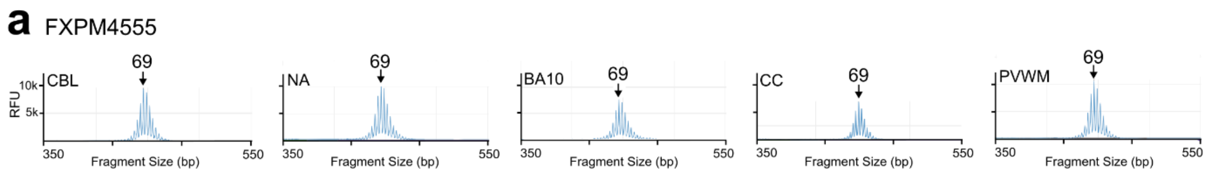


**Supplemental Figure 3. Intra-individual *FMR1* TP-PCR in low (ii) individual.** All molecular characterization is determined from case FXPM 4555. Region assessed: cerebellum (CBL), nucleus accumebens (NA), BA10, corpus callosum (CC), and periventricular white matter (PVWM). Premutation allele detected at 69 repeats in all regions.

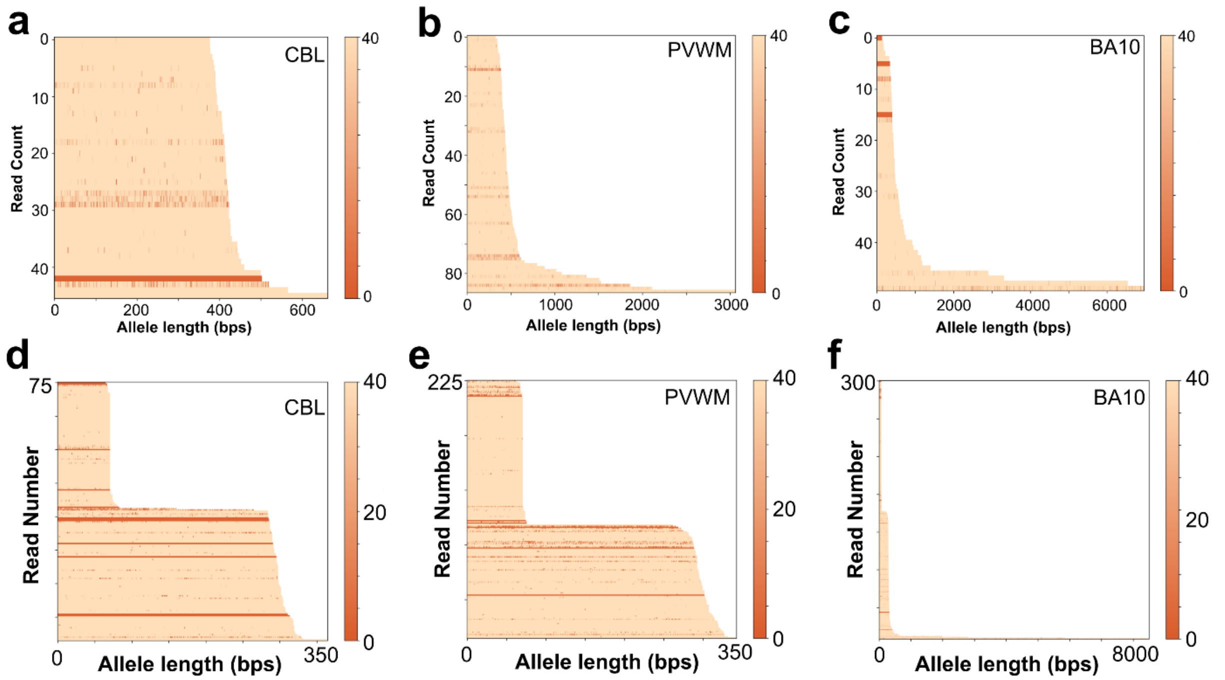


**Supplemental Figure 4. Intra-individual patterns of *FMR1* methylation in FXTAS. a-c)** Base-call quality plots for cerebellum (a), periventricular white matter (b), BA10 (c) for male FXPM 5006. Base call quality scale is 0 = poor, 40 = high quality. **d-f)** Base-call quality plots for cerebellum (a), periventricular white matter (b), BA10 (c) for male FXPM 1008-20-RF.

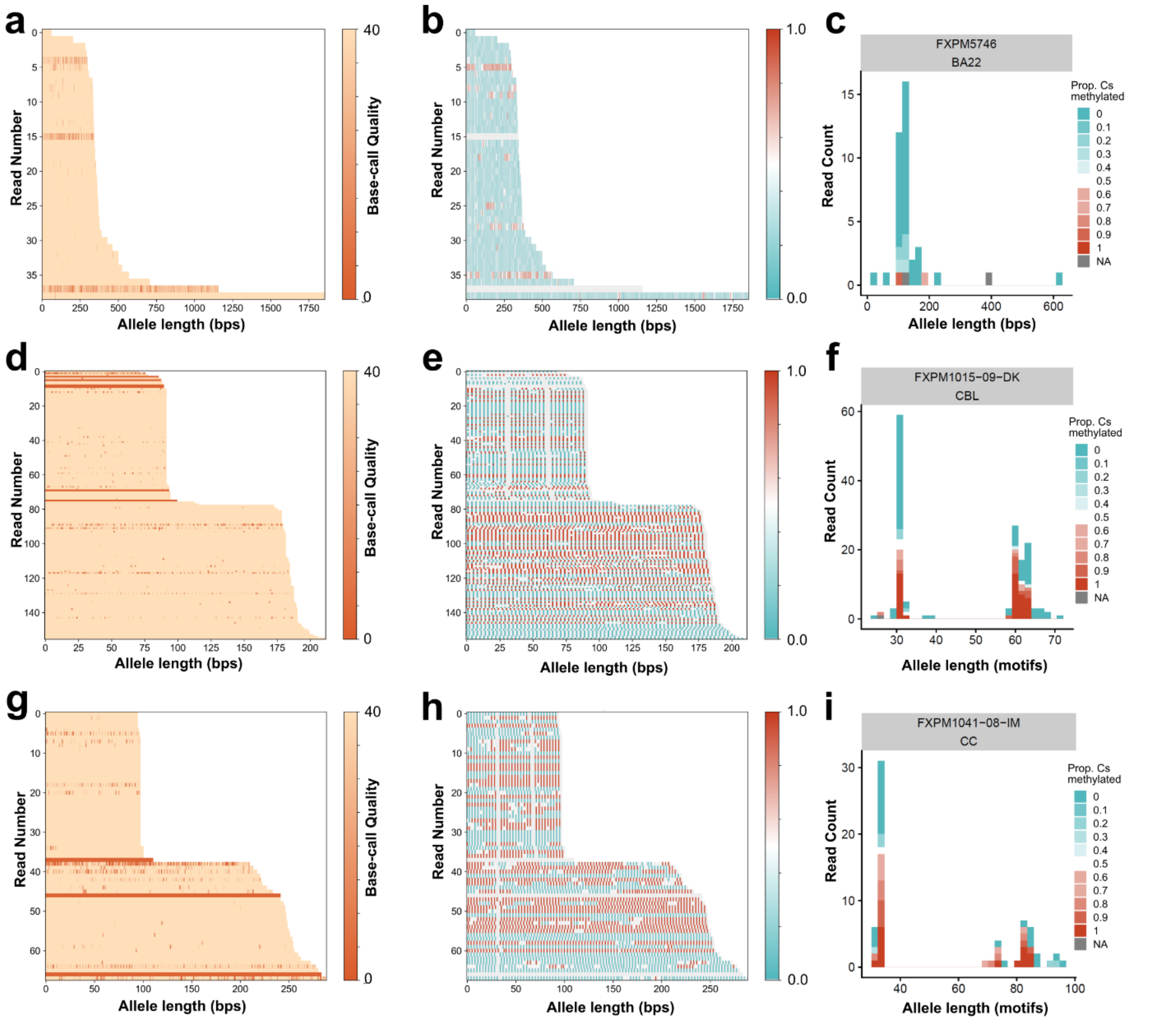


**Supplemental Figure 5. Patterns of *FMR1* methylation in a larger FXTAS cohort. a-c)** Base-call quality plot (a), waterfall methylation plot (b), and allele distribution plot (c) for BA22 of male FXPM 5746. **d-f)** Base-call quality plot (d), waterfall methylation plot (e), and allele distribution plot (f) for cerebellum of female FXPM 1015-09-DK. **g-i)** Base-call quality plot (g), waterfall methylation plot (h), and allele distribution plot (i) for corpus callosum (CC) of female FXPM 1041-08-IM.

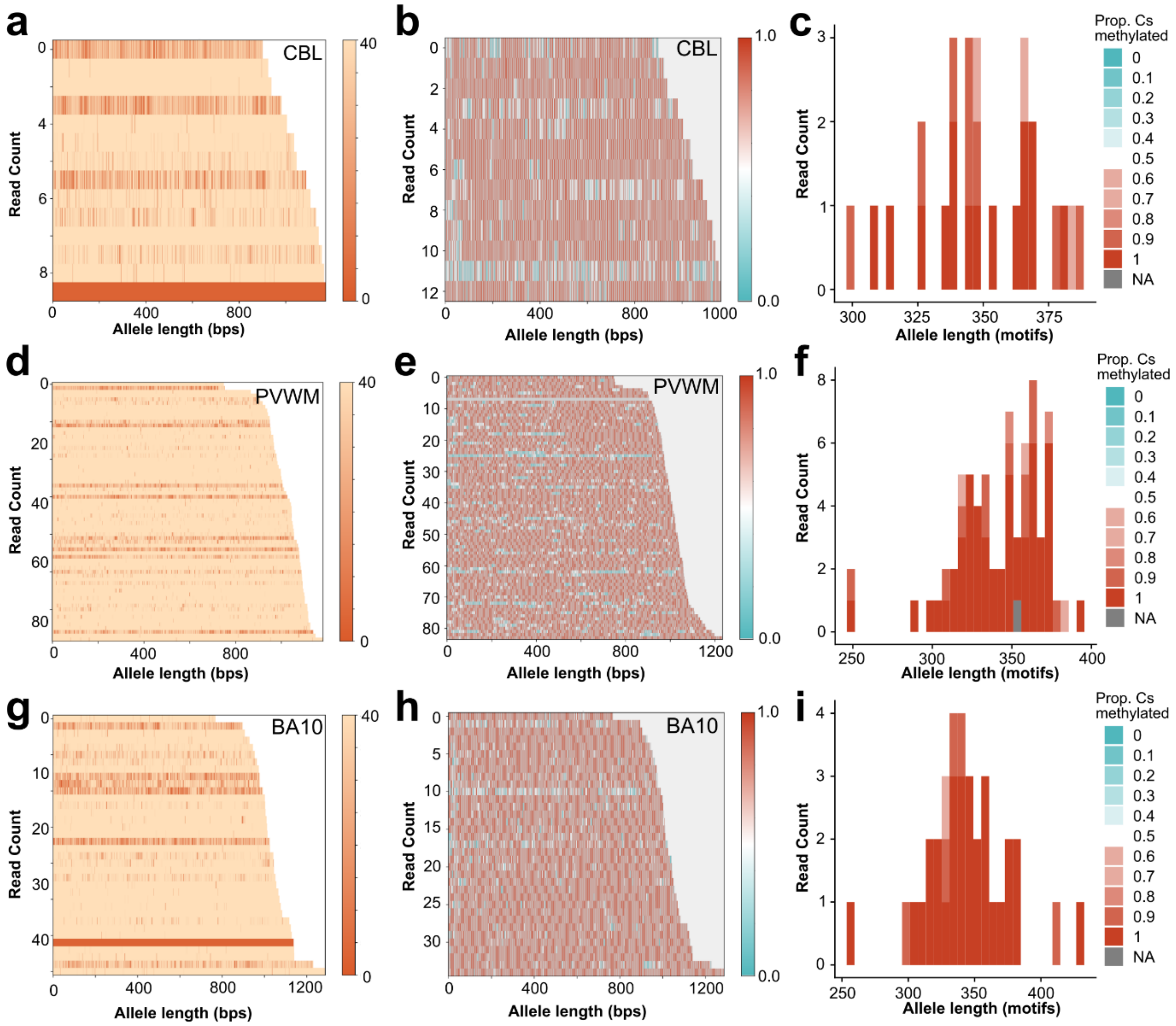


**Supplemental Figure 6. Intra-individual patterns of *FMR1* methylation in FXS case.** All plots are from case FXS 5319. **a-c)** Base-call quality plot (a), waterfall methylation plot (b), and allele distribution plot (c) for cerebellum (CBL). **d-f)** Base-call quality plot (d), waterfall methylation plot (e), and allele distribution plot (f) for periventricular white matter (PVWM). **g-i)** Base-call quality plot (g), waterfall methylation plot (h), and allele distribution plot (i) for BA10.

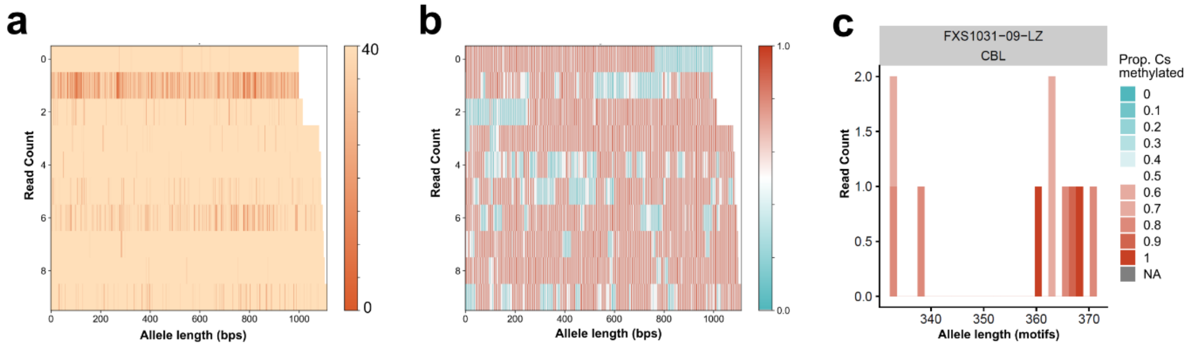


**Supplemental Figure 7. Hypermethylation of *FMR1* is common in FXS. a-c)** Base-call quality plot (a), waterfall methylation plot (b), and allele distribution plot (c) for cerebellum of male FXS 1031-09-LZ.
